# Repurposing Old Antibodies for New Diseases by Exploiting Cross Reactivity and Multicolored Nanoparticles

**DOI:** 10.1101/2020.03.17.995738

**Authors:** Cristina Rodriguez-Quijada, Jose Gomez-Marquez, Kimberly Hamad-Schifferli

## Abstract

We exploit the cross-reactivity of dengue (DENV) and zika (ZIKV) virus polyclonal antibodies for nonstructural protein 1 (NS1) to construct a selective sensor that can detect yellow fever virus (YFV) NS1 in a manner similar to chemical olfaction. DENV and ZIKV antibodies were screened for their ability to bind to DENV, ZIKV, and YFV NS1 by ELISA and in pairs in paper immunoassays. A strategic arrangement of antibodies immobilized on paper and conjugated to different colored gold NPs was used to distinguish the three biomarkers. Machine learning of test area RGB values showed that with two spots, readout accuracies of 100% and 87% were obtained for both pure NS1 and DENV/YFV mixtures, respectively. Additional image pre-processing allowed differentiation between all 4 DENV serotypes with 92% accuracy. The technique was extended to hack a commercial DENV test to detect YFV and ZIKV by augmentation with DENV and ZIKV polyclonal antibodies.

**TOC GRAPHIC:** 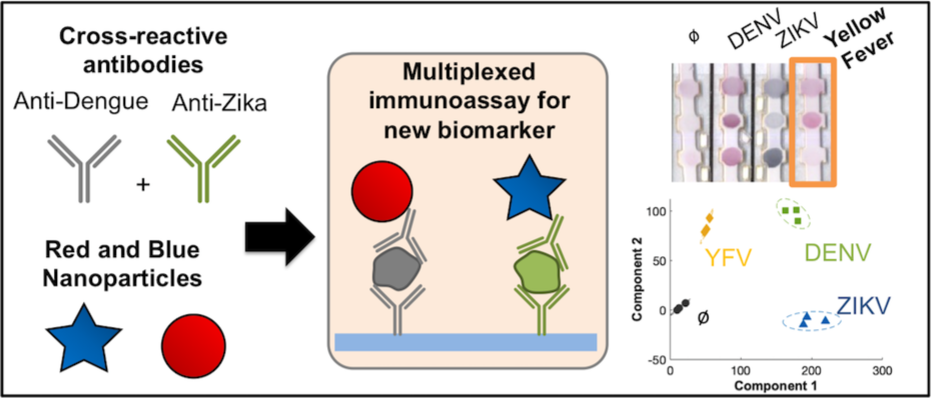

---

Diagnostic tools are key for responding to infectious disease outbreaks, because they provide critical information for patient care, resource allocation, disease containment, and public health surveillance. Point-of care (POC) diagnostics have gained attention for emergency situations because they are inexpensive, portable, operable by non-experts, and deliver results within minutes. ^1, 2^ The most widely used POC diagnostics are paper-based lateral flow immunoassays, where a biological fluid is added to a paper strip. The fluid wicks through the strip by capillary action,^3-5^ and two colored lines appear for a positive test, and one line for negative, which can be read out by eye. This color is due to the presence of gold nanoparticles (NPs) conjugated to the antibody raised against the target biomarker. ^6^ Paper-based immunoassays are valuable diagnostics in resource-limited settings.^7^ While lab-based assays like PCR and ELISA have higher sensitivity and specificity, they are difficult to implement in fieldable, decentralized situations because they require electricity, lab infrastructure, experts to operate them, and cold chains for required reagents.

With every new epidemic, diagnosis is essential during the initial outbreak period for disease containment and surveillance. Unfortunately, for new outbreaks, the necessary biological reagents for diagnosis are not yet available, and an explosive rise in infections can leave clinicians with limited recourse.^8^ Generating new antibodies requires an enormous amount of time and money, where the entire process from selection to manufacturing takes 16-24 months with costs reaching $100M.^9, 10^ In the 2014 Ebola and 2015 Zika outbreaks, even with concerted multi-agency efforts to accelerate the development of paper-based rapid diagnostics, production of a rapid test took 11-14 months, arriving long after the initial stages of both outbreaks. Clearly, the lack of a rapid antibody production method impedes response to epidemics that spread at accelerated rates when it is most crucial.^11^

However, an opportunity lies in the strategic use of antibody cross-reactivity. If cross-reactive antibodies can be used to detect the biomarker of the emerging outbreak, this creates the possibility of leveraging antibodies that have already been mass-produced, effectively bypassing production times. Antibody cross-reactivity is common, such as in commercial diagnostics which can sometimes exhibit false positives for similar diseases, *e.g*. dengue tests showing false positives for zika.^12-15^ Thus, this indicates that the test can be used to detect zika. However, cross-reactivity has the obvious drawback that the two diseases cannot be distinguished from one another, preventing differential diagnosis. This is potentially dangerous for diseases that co-circulate and share vectors and symptoms. For example, zika, dengue, chikungunya, and yellow fever, are all transmitted by the same mosquito vector, and possess similar initial symptoms but drastically different disease outcomes. Thus, leveraging cross-reactivity must be done in a way where one can still distinguish the emerging virus from the original one against which the antibodies were raised.

One solution is to use antibodies as *selective* sensors as opposed to specific ones. While specific sensing relies on an agent that binds only to the target and nothing else, selective sensing uses an array of agents in combination with pattern recognition to distinguish between multiple species.^16^ Chemical olfaction, a form of selective sensing, is a powerful approach that can distinguish between hundreds of highly similar small molecules.^17, 18^ Olfaction relies on multiple agents to bind to analytes with a range of specificities, where some are highly specific, and others more promiscuous. Here, we show that selective sensing can be adapted to the antibody-antigen interaction, enabling discrimination between multiple antigens using cross-reactive antibodies. Furthermore, a selective approach has the potential to leverage cross-reactive antibodies that may be on hand, thus taking advantage of existing stockpiles.^19^ Typically, olfaction uses an array of affinity agents to effect a colorimetric change when the analytes are present. Sandwich immunoassays, however, require a double binding event to produce a signal. We introduce a variable signal *via* the NP-antibody (Ab) conjugate by utilizing the NP optical properties. The color of gold NPs can be tuned by changing their size or shape,^20^ producing different test line colors.^21^

We show that paper-based assays can detect viral antigens in a manner similar to chemical olfaction by exploiting cross-reactivity and NP-antibody conjugates of different NP colors. Using only polyclonal antibodies raised against dengue virus (DENV) serotype 2 and Zika virus (ZIKV) NS1, the assay detected yellow fever virus (YFV) while still distinguishing all 4 serotypes of DENV and ZIKV by exhibiting different colorimetric patterns. Machine learning trained the assay with test line RGB values to enable biomarker discrimination. Color deconvolution with pseudocolors improved classification of biomarkers with the highest sequence similarity. We extend the technique to hack an off-the-shelf dengue diagnostic by augmenting it with DENV and ZIKV antibodies and blue NPs so it can detect DENV, ZIKV, and YFV. We show that with different colored NPs and antibodies with a range of cross-reactivities, one can construct a YFV assay entirely out of antibodies not raised specifically against YFV.

## RESULTS/DISCUSSION

The assay target was NS1, a common biomarker for flaviviruses^22-25^ because it is present at high levels in patients soon after infection and before IgG and IgMs are generated. It allows for early detection, as early as the first day of disease onset when patients exhibit nonspecific symptoms. While YFV outbreaks have been occurring for many years and has a vaccine, new emergences are still occurring, including one in Brazil in 2018. Currently, there is no YFV rapid test similar to the NS1 tests for DENV or ZIKV.^26^

### Multicolor nanoparticles

Star-shaped gold NPs, or gold nanostars (GNS) have a surface plasmon resonance (SPR) peak which shifts through the optical spectrum with GNS arm length, resulting in different colors.^27-29^ To evaluate which NP colors would provide the best color separability, we quantitatively analyzed the RGB differences between the GNS and spherical red NPs for color differentiation.^30^ GNS were synthesized with varying conditions (Methods/Experimental) to result in NPs of different colors. We evaluated the colors of the NPs spotted onto nitrocellulose (**Figure 1a)**, as the final assay would compare colors on paper. NPs exhibited different RGB values (**Figure 1b**). To quantify their differences in RGB space, we calculated the Euclidean distance between the NP RGB values and compared them in a matrix, normalizing them to 1 for the highest value (**Figure 1c**). Values of 0 (heat map, green) represented no Euclidean distance and thus the highest color overlap between two samples, while values approaching 1 (heat map, yellow) represented the highest Euclidean distances and thus lowest overlap. Values of 1 thus indicated higher probability of color discrimination. The best separation was determined to be for red NPs and blue GNS450 (a Euclidean distance value of 1.0). GNS spectra exhibited an SPR peak at 630 nm (**Figure 1d, blue**). GNS450 were characterized for their hydrodynamic size (*D*_*H*_) by dynamic light scattering (DLS, **Figure 1e**) and core size by TEM (**Figure 1f**). Zeta potential measurements confirmed a negative charge (**Figure 1g**). Purchased red spherical NPs 40 ± 4 nm in diameter possessed an absorption peak at 527 nm (**Figure 1d, red**).

**Figure 1.**
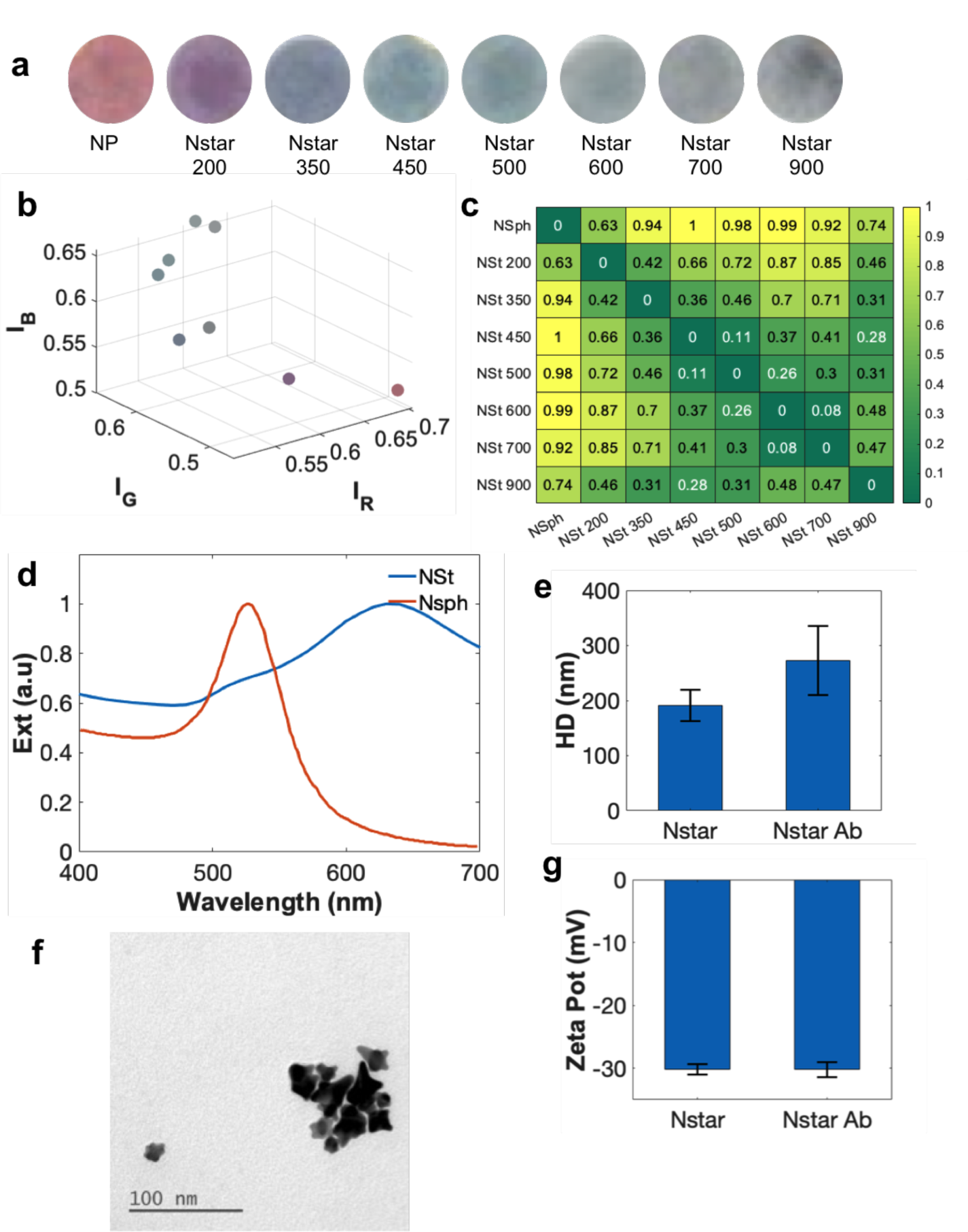
NP synthesis and characterization. a) images of spotted NP and GNS samples, b) RGB values of spotted NP and GNS, c) Similarity matrix of the RGB values of the spots. The Euclidean distance of the RGB values of each of the spotted GNS samples were calculated using [RGB(GNS_1_)-RGB(GNS_2_)]^2^ and normalized from 0 to 1, for the minimum and maximum distances, respectively. A heatmap was generated from these distances, where yellow corresponds to the maximum distance of 1 and 0 for the minimum distance. d) UV-vis of the optimal particles, NP and GNS450. The SPR peak at 630 nm was used to quantify GNS450 concentration using a molar extinction coefficient of ε = 4.3 × 10^8^ M^-1^ cm^-1^ e) DLS of GNS450 with *D*_*H*_ of 191.3 ± 28 nm and GNS450-pDENV,. f) TEM of GNS450 showed an internal diameter of 16.0 ± 1.5 nm, g) zeta potential measurements of GNS450 and GNS450-pDENV (Red and Blue).

### Antibody library

A set of DENV and ZIKV antibodies from commercial sources were screened for their ability to bind to NS1 from DENV, ZIKV, and YFV. The polyclonal DENV (pDENV) was raised against DENV NS1 serotype 2. To investigate cross-reactivity, we divided the DENV2 and ZIKV NS1 sequences into 23 random segments 15 residues in length to represent the potential binding epitopes. A Basic Local Alignment Search Tool for proteins (BLASTP) was performed against the whole DENV1-4, ZIKV and YFV NS1. NS1 sequence varies among dengue serotypes with a sequence similarity of ∼75%, and has a 61% similarity with ZIKV and 56% with YFV ^31^ (**Figure 2a, Supporting Info, Figure S1**). NS1 for DENV2 was used in the comparison as that was the serotype used for polyclonal antibody generation.

**Figure 2.**
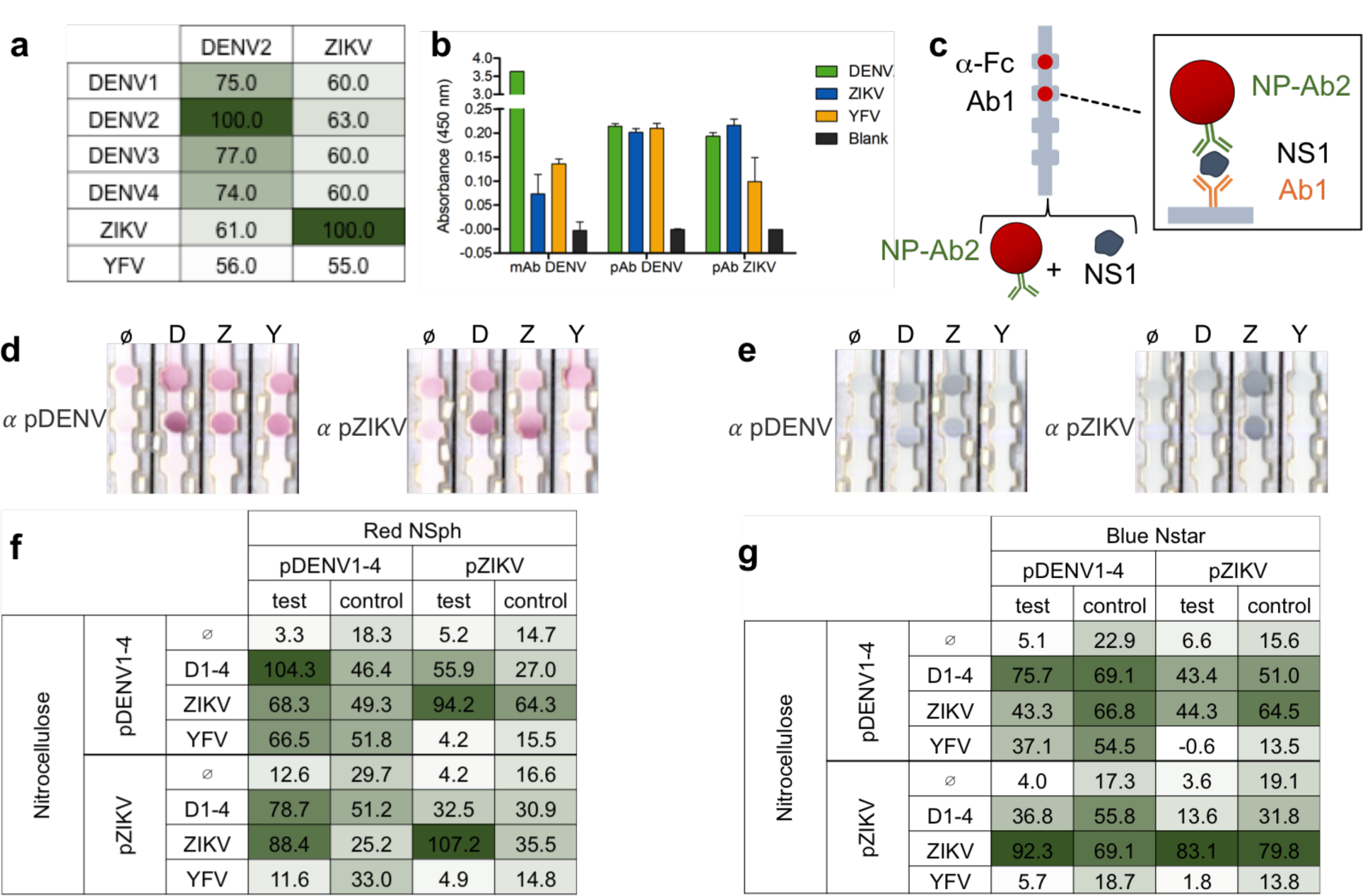
Selection of the antibody panel for repurposing. a) sequence identity between DENV 2 and ZIKV and DENV1-4, ZIKV, and YFV in %. b) ELISA of antibodies in the panel binding to pan DENV NS1 (green), ZIKV NS1 (blue), YFV NS1 (yellow), and the control of no NS1 (black). c) schematic of antibody pair screening where strips had an antibody immobilized at the test line (Ab1) and anti-Fc at the control area. Strips were run by immersing them in a solution containing the NPs or GNS conjugated to different antibodies (Ab2) and NS1 of DENV, ZIKV, and YFV. d) examples of strips run with red NP-pDENV and e) GNS-pZIKV NS1 and DENV, ZIKV, and YFV with immobilized pDENV (left) and pZIKV (right). Test line intensities and heat map of all the pair screenings run with f) red NPs and g) blue GNS. Selected pairs for the assay circled in red.

ELISA was used to probe antibody binding to NS1 targets. Both polyclonal antibodies bound to NS1 of pan (1-4) DENV (green), ZIKV (blue), while exhibiting differences for YFV (yellow, **Figure 2b**). Controls of no NS1 (black) were run. The monoclonal DENV antibody (mDENV) bound strongly to DENV NS1, but less to ZIKV and YFV.

The antibodies were then screened for their abilities to bind to DENV, ZIKV, and YFV NS1 as pairs in dipstick immunoassays. ^31-33^ Each antibody was conjugated to both the red NPs and the blue GNS, and also immobilized on nitrocellulose strips at the test line. Anti-Fc IgG antibodies were immobilized on the control area. The strips were run with each of the NS1 biomarkers. The bottom of the strip was submersed in the solution of NP-Ab + NS1, which wicked up the strip by capillary action. Appearance of color at the test area indicated that the antibodies could bind to the NS1 as a pair, accumulating NPs (**Figure 2c**). Color at the control area indicated that the NP-Abs bound to the immobilized anti-Fc, verifying proper fluid flow. Example strips for red NPs conjugated to polyclonal anti-DENV (NP-pDENV) and blue GNS conjugated to polyclonal anti-ZIKV (GNS-pZIKV) run with immobilized pDENV and pZIKV (**Figure 2d,e**) are shown.

A heat map was created based on the test line intensity values for each antibody pair, where greater intensity in the table indicated higher test area intensity (red NP-Ab, **Figure 2f**, blue GNS-Ab, **Figure 2g)**. The degree of binding of the pairs varied for each biomarker and also with NP shape, providing a range of binding strengths. More importantly, YFV NS1 could be recognized by some antibody pairs but not all, demonstrating that the panel possessed differential cross-reactivity. Interestingly, mDENV did not bind to YFV or DENV1-4 NS1 in pairs (**Supporting Info, Figure S2**), despite being able to bind to DENV NS1 in ELISA. This could be due to the epitope being blocked in a pair format. Thus, mDENV was not used for assay designs.

We quantified detection limits (LODs) of the biomarkers for the selected antibody pairs. NS1 concentration was varied and the resulting test area intensity was measured (**Figure 3a**). Test area intensities of red NP-Ab (red squares, **Figures 3b-g**) and blue GNS-Ab (blue stars, **Figures 3b-g**) with immobilized pDENV and pZIKV exhibited Langmuir-like curves, increasing with NS1 and then saturating.^34^ Fits to a modified Langmuir isotherm (red and blue lines, **Figure 3b-g**) ^35^ allowed calculation of the LODs, which ranged from 1.8 - 972.4 ng/mL (**Table 1**). The YFV LOD was below reported values for positive samples (178-4600 ng/mL), supporting its utility in diagnostic assays.^36^

**Table 1.** Limits of Detection (LODs) of dipstick assays of pairs in ng/ml.

|  | Immobilized pDENV |  | Immobilized pZIKV |  |
| --- | --- | --- | --- | --- |
|  | NP | GNS | NP | GNS |
| DENV NS1 | 1.8 | 5.8 | 116.3 | 972.4 |
| ZIKV NS1 | 6.1 | 42.8 | 44.0 | 20.5 |
| YFV NS1 | 24.7 | 101.3 | 0.0 | 0.0 |

**Figure 3.**
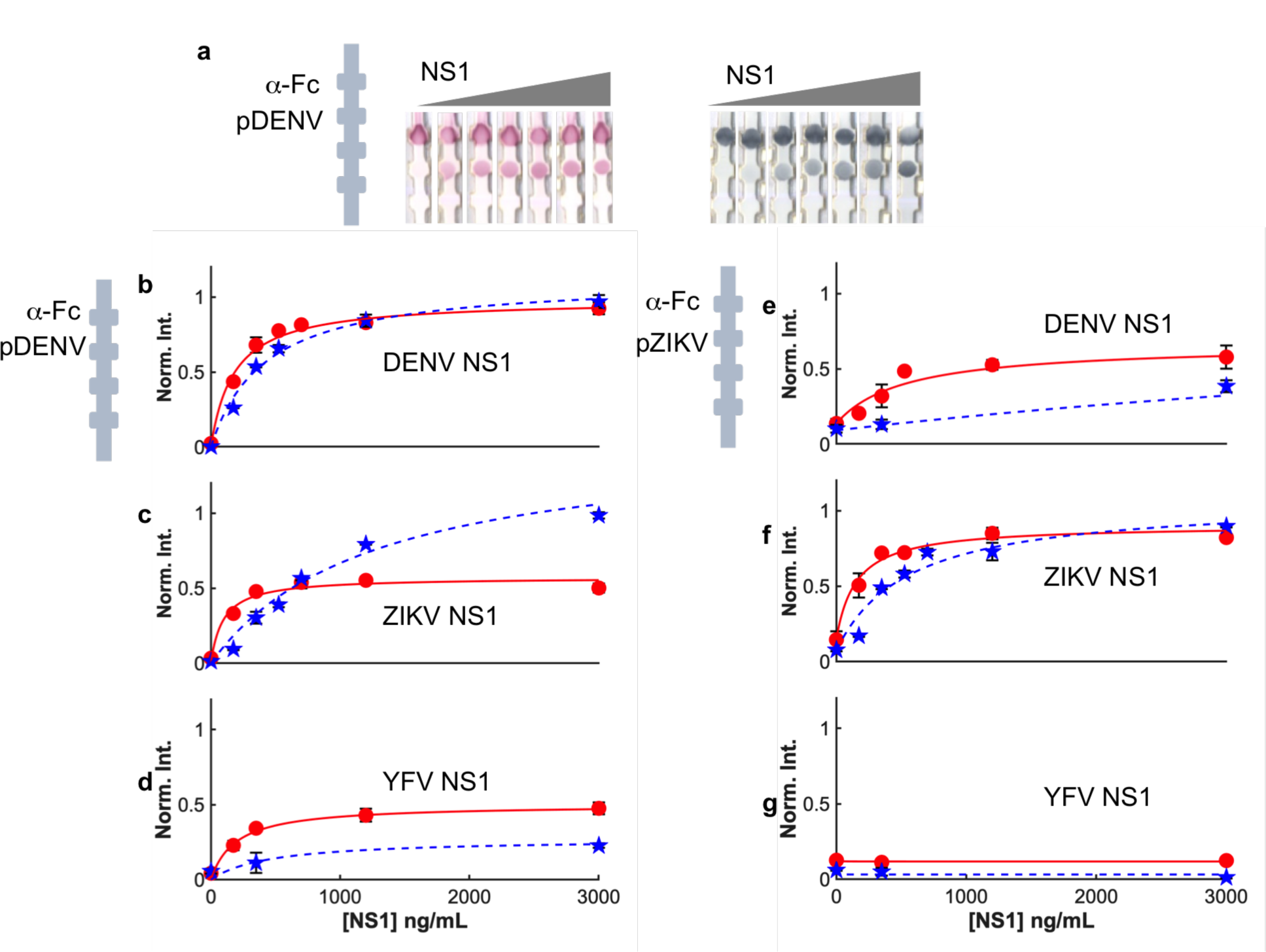
Titration curves of selected antibody pairs. a) Images of strips with immobilized pDENV run with red NP-pDENV and increasing DENV NS1 concentration (left) and blue GNS-pZIKV with increasing ZIKV NS1 concentration (right). Test line intensities of immobilized pDENV strips run with red NP-pDENV (red circles) and blue GNS-pZIKV (blue stars) and b) DENV NS1, c) ZIKV NS1, and d) YFV NS1. Test line intensities of immobilized pZIKV strips run with red NP-pDENV (red circles) and blue GNS-pZIKV (blue stars) and c) DENV NS1, d) ZIKV NS1, and e) YFV NS1. LODs calculated as the minimum concentration that gives a difference in intensity between the control and test lines 3X larger than the standard deviation of the control intensity.

Most notably, titration curve intensities varied with antibody pair and NS1 identity, suggesting that the pairs can differentiate between the NS1. Red and blue NP-Ab conjugates resulted in different test line intensities, which was most likely due to different antibody coverages on the NPs from differing conjugation efficiencies of both NP shapes.^37^ This variation in intensity indicates that for a given NS1, red and blue contributions differ.

### Olfactory array to distinguish DENV, ZIKV, YFV

Using the information from screening and titration curves, we strategically combined the pDENV and pZIKV antibodies and NP-Ab conjugates to make a DENV/ZIKV/YFV test. A two-spot test was constructed using pZIKV as the capture antibody at one position, and pDENV at the other (**Figure 4a**). pZIKV was conjugated to the blue GNS, and pDENV to the red NPs. Tests were run with mixtures of both NP-Ab conjugates against NS1 from DENV, ZIKV, YFV spiked in human serum (HS). When DENV was the target, DENV NS1 from all four serotypes were mixed together (pan DENV NS1).

**Figure 4.**
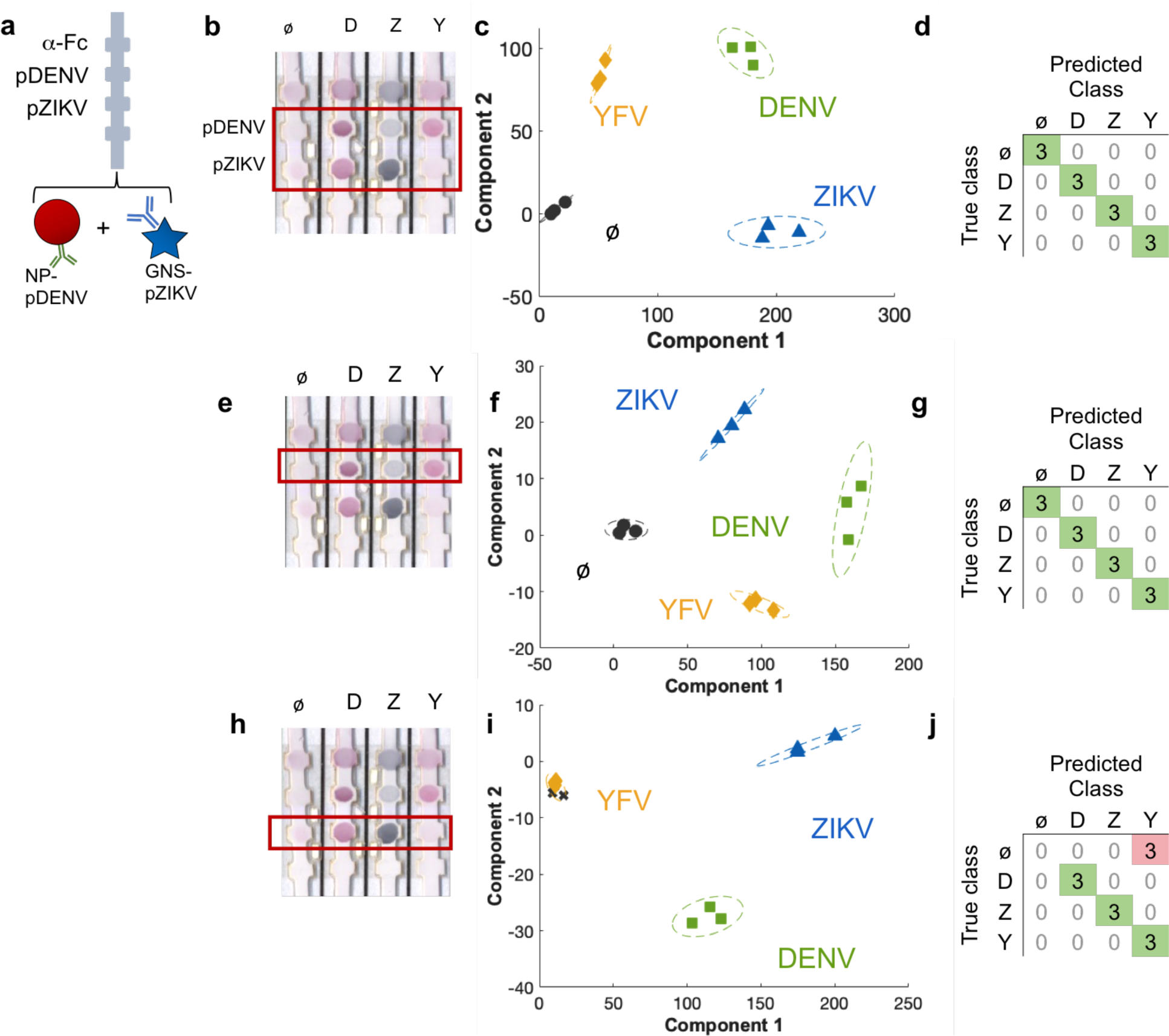
Multiplexed assay that can distinguish ZIKV, DENV, YFV NS1, analyzed in 3 different ways. a) Schematic of the final assay configuration, which is red NP-pDENV and blue GNS-pZIKV run with a strip with immobilized pDENV and ZIKV. b) Images of the assay run with blank (ø), DENV NS1 (D), ZIKV NS1 (blue), and YFV NS1 (Y). c) PCA analysis of the two test areas boxed in red and clustering shown for the DENV (green squares), ZIKV (blue triangles), YFV (yellow diamonds) NS1 and the blank (black squares), d) confusion matrix of the PCA analysis shows 100 % accuracy between predicted and true classes. Correct predictions: on-diagonal, green; incorrect predictions: off-diagonal, pink. e) Analysis using only the upper test area (immobilized pDENV, red box) and f) PCA analysis and g) resulting confusion matrix with 100 % accuracy. h) Analysis using only the lower test area (immobilized pZIKV, red box) and i) PCA analysis show misclassifications on the blank (black Xs) and j) confusion matrix with 75% accuracy.

Different color patterns resulted when each of the biomarkers were run (**Figure 4b**). Pan DENV NS1 gave rise to spots at both test areas that were redder in color. This suggested that pairs were formed for both immobilized pDENV and pZIKV, and that red NP-pDENV tended to pair better when run with DENV NS1. When run with ZIKV NS1, the assay also resulted in spots bluer in color at both areas. This suggested that both immobilized pDENV and pZIKV tended to pair better with blue GNS-pZIKV in the presence of ZIKV NS1. For YFV NS1, a red spot appeared at the pDENV test area, but the pZIKV test area remained blank. This was corroborated by the fact that pZIKV did not exhibit strong cross-reactivity with YFV NS1 in the antibody pair screening (**Figure 2f,g)**. This shows that a different RGB pattern resulted depending on the biomarker present, enabling discrimination between the biomarkers using only ZIKV and DENV polyclonal antibodies.

To investigate clustering, we quantified the test area colors and performed principal component analysis (PCA) with supervised learning. PCA used the RGB values for each of two test areas (giving rise to 6 components), and was performed (Methods/Experimental) with 10-fold cross validation of the tests run in triplicate (**Figure 4c**, yellow diamonds, YFV; green squares, pan DENV; blue triangles, ZIKV; blank, black circles). LDA was used to build a classification model by feeding the same RGB intensity values at both spots (6 components). A 10-fold cross validation was performed to fit the model trained with the tests run in triplicates. The validation accuracy of the tests was determined by measuring the correctly and incorrectly classified samples by the LDA model, and visually representing classifications in a confusion matrix, where the rows represent the true classes, and columns the predicted classes (**Figure 4d**). Comparing correctly classified samples (diagonals, green) *vs*. misclassified samples (off diagonals, white), showed that PCA could classify with 100% validation accuracy.

Going further, we investigated the ability to distinguish samples using only the upper test spot with immobilized pDENV (**Figure 4e**). PCA was trained against the 3 NS1s and no NS1 using only the upper spot (with 3 components, **Figure 4f**) and could classify samples with 100% accuracy (**Figure 4g**). Even though it is difficult to distinguish between the DENV and YFV test colors by eye, machine learning could distinguish them successfully. When using only the test area with immobilized pZIKV (**Figure 4h**), PCA (**Figure 4i)** correctly classified DENV and ZIKV NS1, but could not distinguish YFV NS1 from the blank (black X’s, **Figure 4i**, pink off-diagonal, confusion matrix, **Figure 4j**), resulting in a 75% accuracy. This shows that the combination of the two spots is optimal as the pattern can be distinguished by eye, but in practice one pDENV spot could suffice.

### Differentiating NS1 mixtures

We investigated the ability to distinguish mixtures of pan DENV and YFV, which could be necessary in a situation with a co-infection, where both biomarkers may be present. While we do not know what the relevant levels are for a co-infection, we investigated the ability of the test to distinguish between mixtures at different ratios. DENV + YFV NS1 were mixed at different ratios and run in the assays. Except for the 0:1 case, resulting patterns were difficult to distinguish by eye (**Figure 5a**). PCA with 10-fold cross-validation could distinguish between the five mixtures (**Figure 5b**) with a validation accuracy of 73% (**Figure 5c**). Validation accuracy improved to 87% when only three class types were used to train the model (DENV, mixtures and YFV). Two of the nine mixtures were misclassified, where a dipstick run with a 1:2 mixture was classified as pure YFV and a 2:1 mixture was classified as pure DENV (**Supporting Info, Figure S3**).

**Figure 5.**
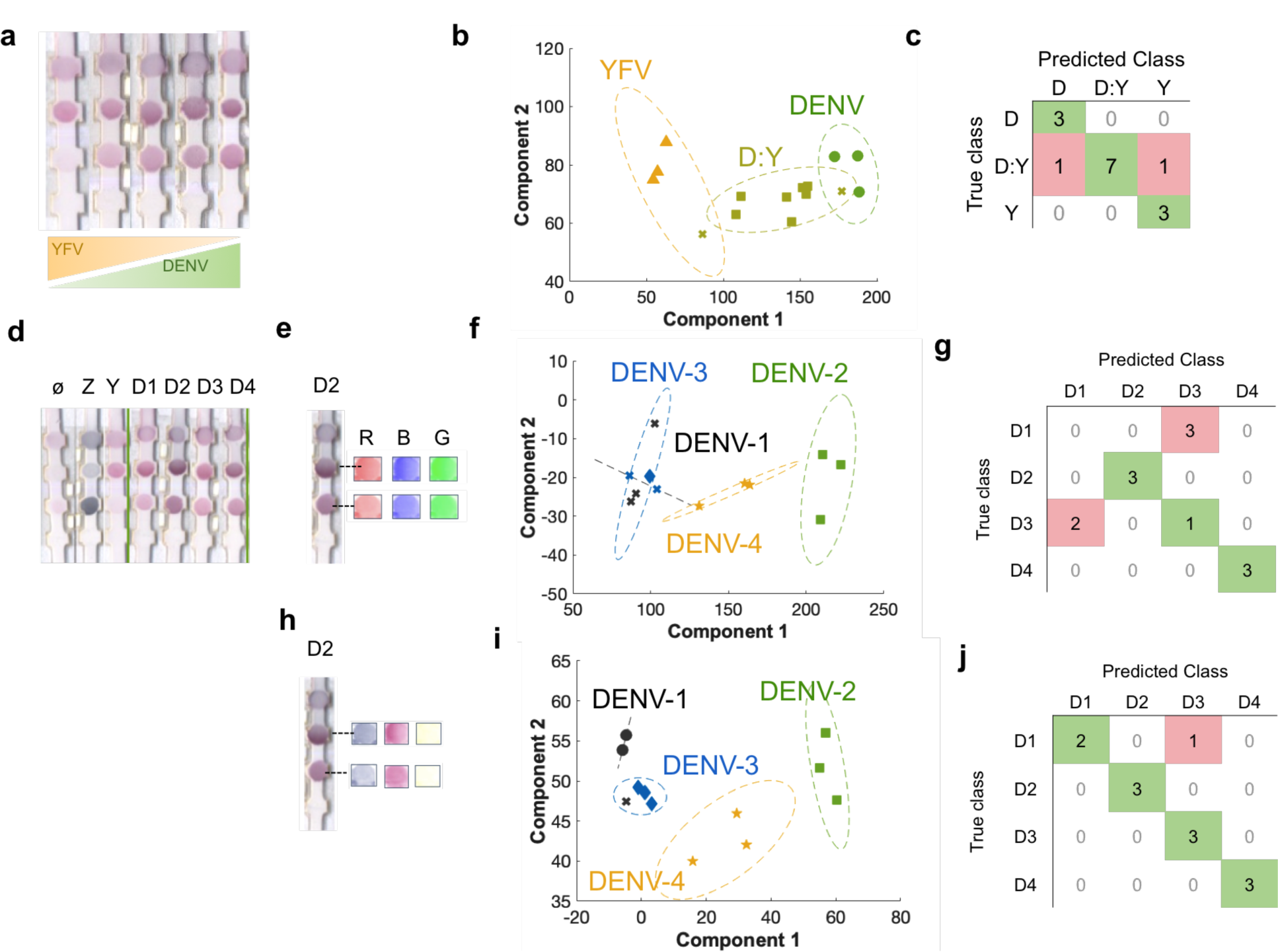
a) Images of test strips run with mixtures of DENV:YFV NS1 at 1:0, 1:2, 1:1, 2:1, 0:1. b) PCA analysis of the different ratio c) Confusion matrix of the 5 classes showing 87 % accuracy. d) Images of strips run with ZIKV (Z), YFV (Y) and DENV 1-4 (D1, D2, D3, D4). e) example strip of DENV-2 test and control areas deconvoluted into R,G,B, channels. f) PCA using RGB values to classify DENV 1 (black), DENV2 (green squares), DENV3 (blue diamonds), and DENV4 (yellow stars). The analysis misclassifies DENV1 and DENV (black and blue X’s), g) confusion matrix of PCA of RGB showing an overall accuracy of 58 %. h) Image deconvolution into pseudocolors representative of the blue GNS, red NP, and everything else. i) PCA analysis of DENV1-4 shows 1 misclassified DENV1 sample, j) confusion matrix with an overall accuracy of 92 %.

### Serotype classification of DENV NS1

DENV has 4 serotypes, and serotype identification is critical for diagnosis,^31^ as secondary infections can lead to dengue hemorrhagic fever and dengue shock syndrome,^38, 39^ so we investigated the assay’s ability to distinguish serotypes. DENV1-4 NS1 run in the assays resulted in red spots at both test areas, which varied in color depending on serotype (**Figure 5d**). PCA of the test area colors could correctly classify DENV2 and DENV4 (green squares, yellow stars, **Figure 5f**). However, DENV1 and DENV3, which are closer in the flavivirus phylogenetic tree, (black and blue X’s) were not correctly classified, resulting in a 58% validation accuracy (**Figure 5g**). ^40, 41^

Although NP and GNS colloidal solutions appeared to be pure red and blue, RGB values of their spots on nitrocellulose were not pure colors, and overlap in RGB space (**Figure 5e**).^42^ This overlap compromises the ability to use RGB to distinguish between nanoparticles, and contributes to a decrease in classification accuracy because a pixel of a given color cannot be accurately separated into how much contribution is from the red NP *vs*. the blue GNS. Color deconvolution is used in histology when stains have RGB overlap, and can be used to separate combinations of stains when they are spatially colocalized. It is an orthonormal transformation of the optical density of a pixel into channels for each stain. To use this in machine learning, first the pseudocolors of the red NPs and blue GNS were identified individually (Supporting Information). This allowed us to determine the contribution of each NP more accurately than in RGB analysis. Then, these three channels were used to train the PCA model: the pseudocolor corresponding to red NPs, the pseudocolor corresponding to blue NPs, and those contributing to neither, or background (**Figure 5h)**. ^43^ By training the model with pseudocolors instead of RGB values on both the test and control areas, validation accuracy increased to 92% (**Figure 5i,j**), which we believe is due to the fact that each pixel can be more accurately assigned to be a red NP or a blue GNS.

### Hacking a commercial DENV test to detect YFV

To extend the capabilities of selective sensing, we hacked a commercial dengue NS1 test (BioCan Dengue NS1) with repurposed DENV and ZIKV antibodies so that it could detect YFV. The Biocan test was not cross-reactive, as it resulted in a positive test for DENV NS1 but negative for ZIKV and YFV (**Figure 6a**). To convert it into a diagnostic that can also detect YFV, the range of cross-reactivities needs to be fully complemented. Following the same design rationale in the YFV tests, we added a cross-reactive antibody pair (**Figure 6b**). pDENV was spotted on the BioCan test strip. Blue GNS conjugated to pZIKV and red NPs conjugated to pDENV were added to the sample and run in the test. When run with DENV, YFV, or ZIKV NS1, the hacked tests exhibited different RGB patterns for the spot and test line (**Figure 6c**). PCA (**Figure 6d**) could distinguish between the different patterns with 100% accuracy (**Figure 6e**). Thus, a DENV diagnostic can be augmented with DENV and ZIKV antibodies to detect YFV, even with no knowledge of the antibody identity in the diagnostic.

**Figure 6.**
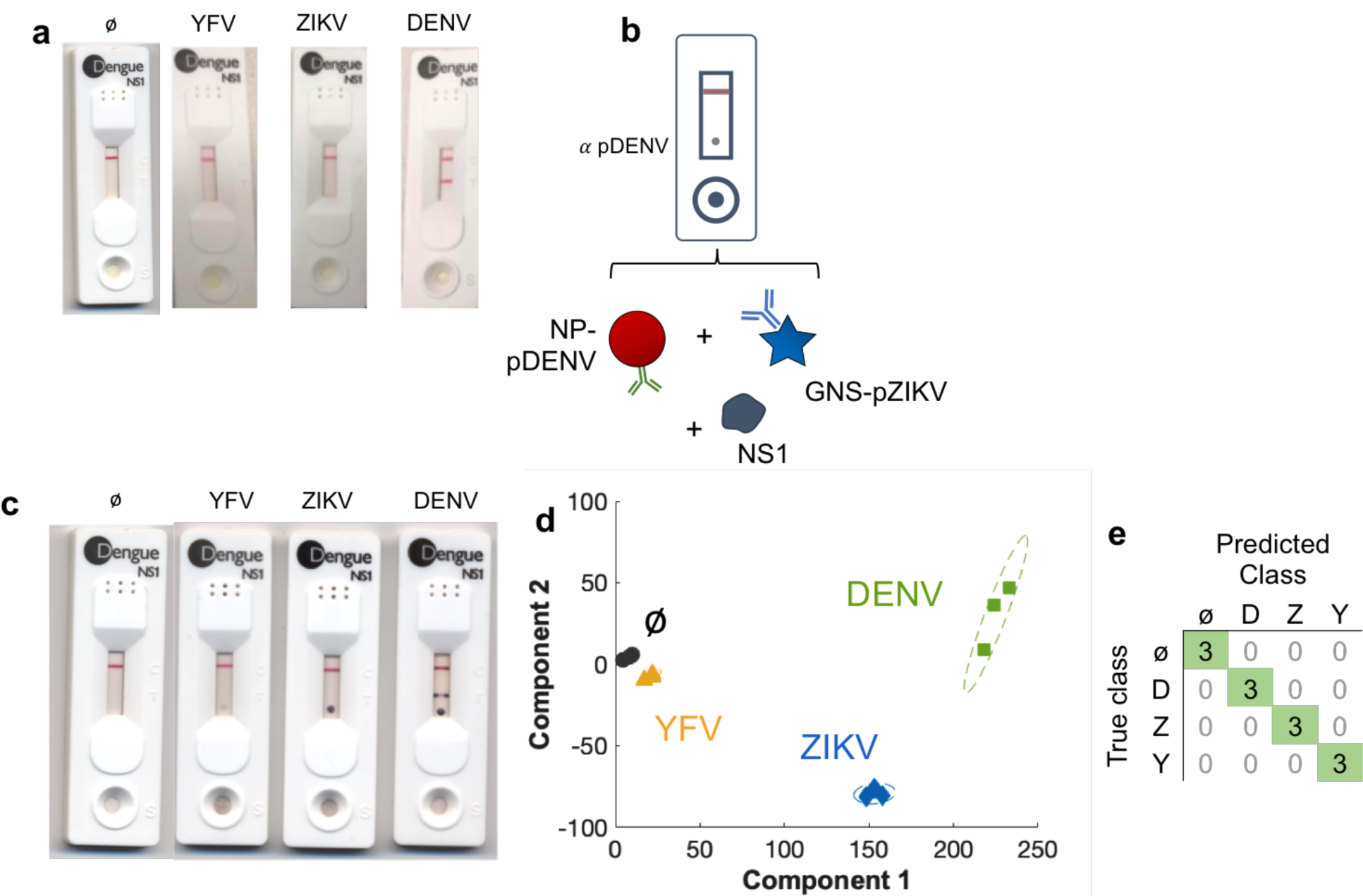
Hacking a commercial DENV test to detect YFV and ZIKV. a) Biocan tests run with NS1 from YFV, ZIKV, and DENV and no NS1 b) configuration of a modified test, where pDENV is spotted down on the strip and the test is run with the addition of NP-pDENV and GNS-pZIKV. c) Hacked Biocan test run with NS1 from YFV, ZIKV, and DENV and no NS1 results in different patterns of the commercial test line and additional spotted area. D) PCA analysis of the Biocan tests for DENV (green squares), ZIKV (blue diamonds), and YFV (yellow triangles, and no NS1 (black circles) and e) confusion matrix with 100 % accuracy.

## CONCLUSION

The unique physical properties of NPs have facilitated the design of selective and olfactory sensors, enabling differentiation of complex mixtures.^44, 45^ The elegantly simple system of a NP modified with only a single surface ligand and three different adsorbed proteins can distinguish subtle differences between multiple cell types or cell phenotypes.^46^ Typically, antibodies are viewed as the canonical specific sensors because they are generated by an immune response specific to an antigen challenge.^16^ Consequently, sandwich immunoassays are used predominantly as single channel devices. However, by extending olfaction to antibodies, which is based on a range of cross-reactivities,^17^ it allows re-purposing of antibodies for a new antigen. Furthermore, we show that commercial diagnostics can be hacked to detect a different antigen for which it was not originally designed.

Cross-reactivity in antibodies is predominantly viewed as undesirable, where cross-reactive antibodies that bind to closely related species are discarded in the antibody selection process. Furthermore, polyclonal antibodies are often perceived as less desirable than monoclonals, but they actually possess distinct advantages for olfaction. Because polyclonals bind to several epitopes with a range of affinities as opposed to a single epitope, they have a higher propensity for cross-reactivity. Potentially, this could contribute to better PCA clustering and consequently higher accuracy. We show that olfactory sensing with cross-reactive polyclonals can even distinguish between DENV serotypes, which is challenging, time consuming, and expensive to achieve with monoclonal antibodies, especially when they are raised to bind to one serotype and not others.^31^ In comparison, polyclonal antibodies are more cost effective and accessible than monoclonals, as their generation does not require a hybridoma. The entire set of experiments conducted here was achieved using stock antibodies, and the final assay used solely two different polyclonal antibodies with only two different NP colors. One drawback of polyclonal antibodies is that batch variability can change binding affinity, but we believe updating the clustering and training with each new batch would be straightforward. This would involve retraining the system, where the new nanoparticle-antibody conjugate would be run in the tests with known antigens, and the RGB values of the test areas trained in PCA.

Infectious disease outbreaks are occurring with greater frequency, and response to them needs to be dramatically improved. We believe that repurposing antibodies for an emerging disease could generate diagnostics during the initial stages of an epidemic, filling a critical gap in disease response before specific antibodies can be mass-produced.^11, 47^ This would be a shift in how we manage rapid response, where we could be better prepared for the next viral outbreak. Our hope is that the demonstration of this approach can be used as a future tool for future outbreaks for which there are not yet rapid diagnostics. Starting with the new biomarker, an arrangement of stockpiled antibodies with multicolored NPs can be arrived at following the procedure of pair-wise screening and machine learning. The equipment required is a camera such as those on a mobile phone or microcontroller board (e.g., Raspberry Pi, Arduino) and a computer to run image analysis software (e.g. Matlab, R). The time required to deploy a POC diagnostic using this repurposing approach could potentially be within months, much shorter than what is required to conventionally generate monoclonal antibodies. While we demonstrate the principle only for repurposing flaviviruses for a YFV test for NS1, we believe that by showing quantitative analysis of its performance and the limits of the approach (i.e., using RGB values *vs*. color deconvolution) will allow others to extend the technology and develop tests for future outbreaks. This approach could potentially be used for other flaviviruses (e.g. using DENV antibodies to detect ZIKV), as well as other virus types (filoviruses, e.g. using Ebola antibodies to detect Marburg), as long as the target is close to the antigen for which antibodies have been developed. Furthermore, this approach could potentially be applied to detect other protein targets such as IgG/IgM antibodies, or the viral proteins themselves. Those local to epidemic outbreaks could be easily trained in this approach, further expediting POC distribution.

## METHODS/EXPERIMENTAL

### Gold Nanostar synthesis

GNS with different arm lengths were synthesized by reduction of HAuCl_4_ (Sigma Aldrich 99.995% trace metal basis) at pH 7.4 as described elsewhere^27^. The concentration of HEPES (Sigma Aldrich 99.5%) was varied from 28 to 126 mM.

### Antibodies

Monoclonal DENV antibodies were obtained from Native Antigen. Polyclonal DENV and ZIKV antibodies were obtained from CTK Biotech.

### Nanoparticle conjugation with antibodies

GNS were centrifuged at 12000 rcf for 12 min and resuspended in Milli-Q water. Afterwards, the solution was centrifuged again at 12000 rcf for 8 min and the pellet was resuspended in 300 µL of Milli-Q water, 100 µL of 40 mM HEPES at pH 7.7 and 10 µL of the antibody at 1 mg/mL (anti-Pan Dengue NS1 and anti-Zika NS1 polyclonal antibodies from CTK Biotech and anti-Pan Dengue NS1 monoclonal antibody from Native Antigen). The solution was left in the orbital shaker for 45 min at room temperature. A PEG backfill was conducted to fill in unpassivated areas on the NP surface, which helps avoid false positives. 10 µL of mPEG was added to the conjugates (MW 5000 >95 % purity from Nanocs) and left for 15 min in the orbital shaker at room temperature. The nanoparticles were centrifuged for 12 min at 10000 rcf to separate the antibody and PEG in excess. Commercial red nanospheres (InnovaCoat 20OD 40 nm, Expedeon) were conjugated following the protocol provided by the manufacturer.

### Gold nanostar characterization

GNS were imaged under the TEM (FEI Tecnai G2) and the dimensions were determined with ImageJ (Version 2.0.0). The GNS sizes and zeta potential before and after antibody conjugation were determined with a Zetasizer Nano ZS (Malvern Instruments). The absorption spectra of GNS and the commercial gold nanospheres were measured on a Cary 5000 UV-Vis-NIR spectrophotometer.

### ELISA

The indirect ELISA was carried out by immobilizing 0.1 µg NS1 per well (Native Antigen) overnight at room temperature. The primary antibody incubation was performed at room temperature for 2 h. The secondary antibodies anti-mouse and anti-rabbit (Abcam) were diluted to 1:1000 and 1:12000, respectively, and incubated at room temperature for 1 h. The assay was read at 450 nm.

### Running strips with spiked NS1

Nitrocellulose strips (Unistart CN140 from Sartorius) were cut with a laser cutter. Antibodies were immobilized by spotting 0.6 µg on the test and control line areas and then were let to dry completely. Anti-Fc IgG antibodies were immobilized on the control area as verification that fluid flow worked correctly. An absorbent pad attached to the top of the strip served as a fluid sink. The bottom of the strips were placed in a nanoparticle solution with 30 µL of human serum with a spiked NS1 concentration of 1 µg/mL unless indicated. Fluid was allowed to wicked up the strip by capillary action towards the absorbent pad. (GB003 Gel Blot paper, Sigma Aldrich).

### Image analysis and Machine Learning

Strips were scanned and the RGB values were determined with ImageJ (Version 2.0.0). Principal component analysis and color deconvolution scripts were performed using Matlab (Version R2018a). RGB values of the test area spots were used to train the PCA model and to visualize the clusterization of the classes. To train the system, the selected NP-Abs from Figures 2 and 3 (red NP-pDENV, blue GNS-pZIKV) were run together against each of the NS1s (DENV, ZIKV, YFV) and no NS1 (negative controls). If both test spots were used in the analysis, the RGB intensities (I_R_, I_G_, I_B_) channels were measured for both test spots, giving rise to 6 components per strip (I_R_, I_G_, I_B_) for position 1, (I_R_, I_G_, I_B_) for position 2). The PCA model used the singular value decomposition algorithm to detect the two components that best discriminate these classes.

LDA was used to build a classification model by feeding the same RGB intensity values at both spots (6 components). A 10-fold cross validation was performed to fit the model trained with the tests run in triplicates. The validation accuracy of the tests was determined by measuring the correctly and incorrectly classified samples by the LDA model, and visually represented in a confusion matrix, with rows representing the true classes and columns the predicted classes.

### Color deconvolution

Color deconvolution into pseudocolors was achieved by following the procedure of Ruifrok et al.^43^ A detailed description is in the Supporting Information.

## ACKNOWLEDGEMENTS

This work was supported by UMass President’s Office of Technology and Commercialization Ventures. CRQ was supported by a Rafael del Pino Fellowship, the Beacon Student Success Fund, a UMass Boston Goranson Award, and an Oracle Sanofi Fellowship. We thank Laura Balcells Garcia for experimental assistance, and Briana Lino for assistance with the commercial assays.

## AUTHOR CONTRIBUTIONS

CRQ, JGM, and KHS conceived the concepts. CRQ and KHS designed the experiments. CRQ carried out the experiments and performed data analysis. Data interpretation was by KHS and CRQ. KHS and CRQ wrote the manuscript.

## COMPETING FINANCIAL INTERESTS

JGM and KHS have filed a patent application on the concept of the work.

## SUPPORTING INFORMATION

Additional information includes the sequence similarity of DENV, ZIKV, and YFV NS1, monoclonal antibody binding in pairs on dipstick assays, and the performance of the ratios of DENV:YFV mixtures, protocol for the screening of antibody pairs, and information on color deconvolution. This material is available free of charge *via* the Internet at http://pubs.acs.org.

