## Supporting Information for "Repurposing Old Antibodies for New Diseases by Exploiting Cross Reactivity and Multicolored Nanoparticles"

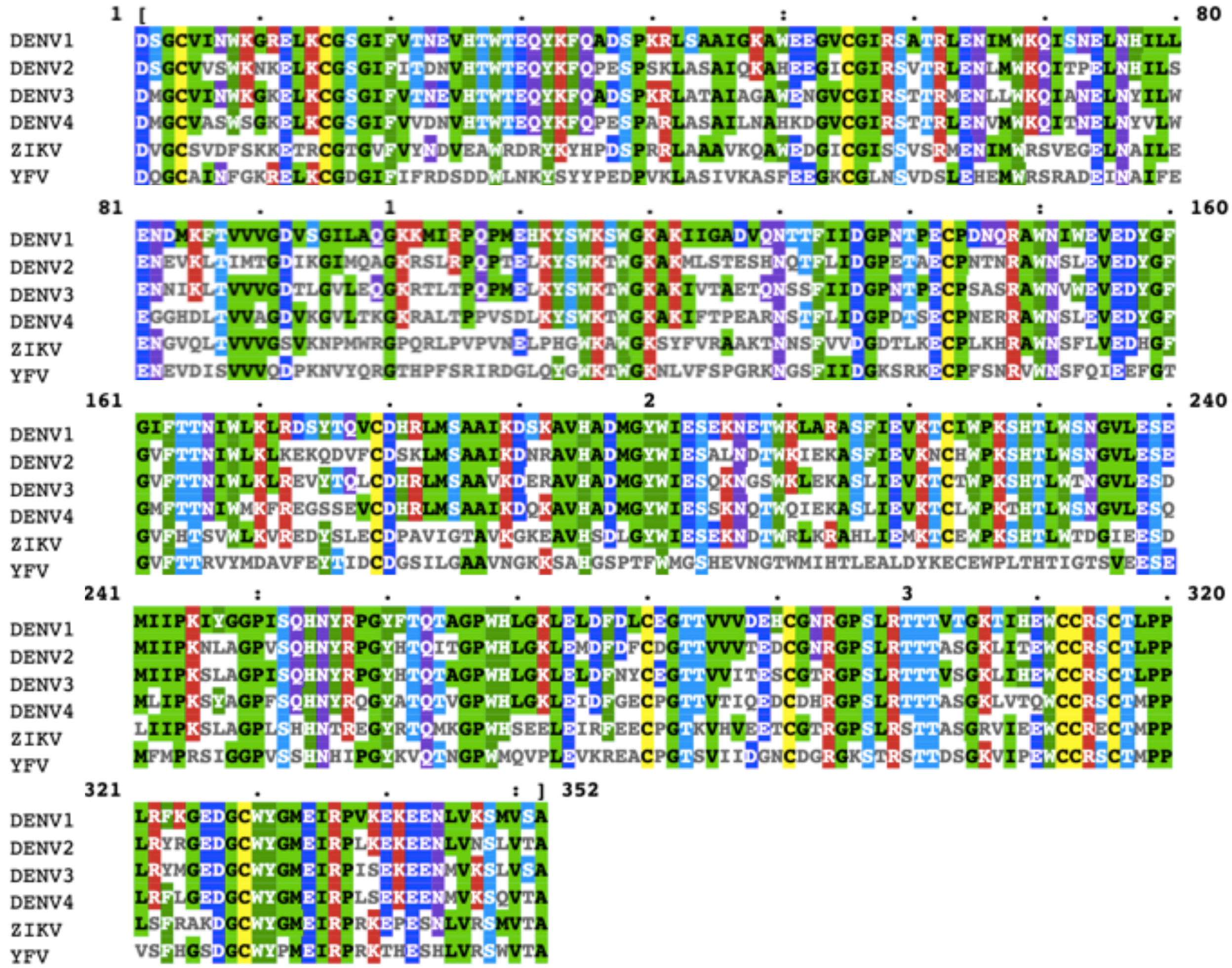

**Figure S1. Sequence similarity between dengue, zika, and yellow fever virus NS1 using protein Basic Local Alignment Search Tool (BLASTp).**

**a**

| | | | Nanoparticle NSph | | |
|---|---|---|---|---|---|
| | | | mAb DENVX | pAb DENVX | pAb ZIKV |
| Nitrocellulose | mAb DENVX | ∅ | | | |
| | | D1 | | | |
| | | D2 | | | |
| | | D3 | | | |
| | | D4 | | | |
| | | YFV | | | |

**b**

| | | | Nanoparticle Nstar | | |
|---|---|---|---|---|---|
| | | | mAb DENVX | pAb DENVX | pAb ZIKV |
| Nitrocellulose | mAb DENVX | ∅ | | | |
| | | D1 | | | |
| | | D2 | | | |
| | | D3 | | | |
| | | D4 | | | |
| | | YFV | | | |

**Figure S2. Immobilized mAb binding to DENV1-4 and YFV NS1 in pairs on a) red NPs and b) blue GNS as a heat map (white, no binding, light green, weak binding).**

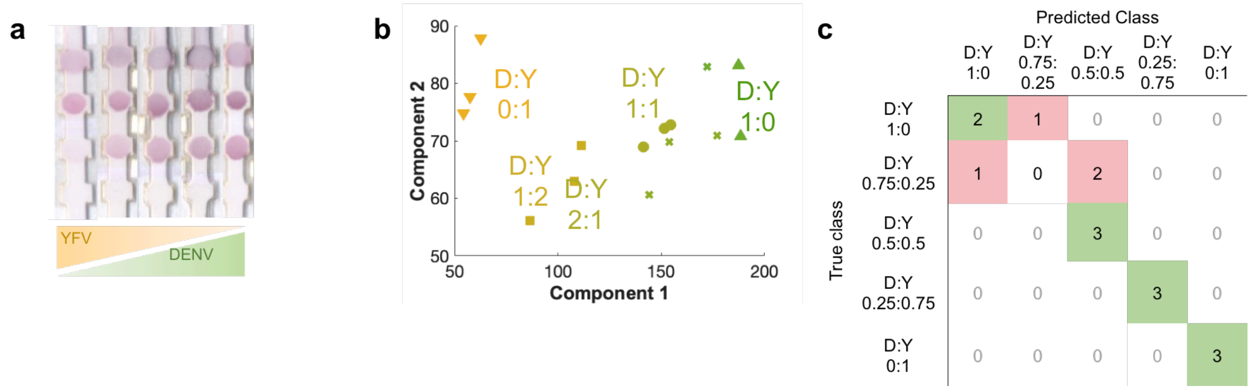


**Figure S3. Mixtures of DENV:YFV NS1 evaluated at different ratios. A) image of strips, b) PCA with the ratios separated out, and c) confusion matrix of the PCA showing accuracy of 73 %.**

### Antibody screening procedure

**For 1st NP color (i.e., red)**
- Vary Ab1 (on nitrocellulose
- Vary Ab 2 (conjugated to nanoparticle)
- Run dipsticks with NS1 for DENV1-4, ZIKV, and YFV and blank

- Image tests for grayscale intensity at test area

- Generate matrix of test line intensities for all combinations

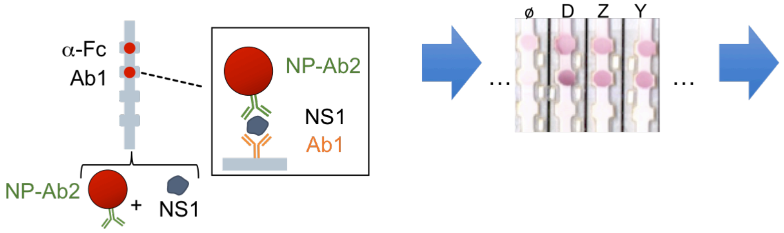


| | | Red NSph | | | |
|---|---|---|---|---|---|
| | | pDENV1-4 | | pZIKV | |
| Nitrocellulose | | test | control | test | control |
| pDENV1-4 | ⌀ | 3.3 | 18.3 | 5.2 | 14.7 |
| pDENV1-4 | D1-4 | 104.3 | 46.4 | 55.9 | 27.0 |
| pDENV1-4 | ZIKV | 68.3 | 49.3 | 94.2 | 64.3 |
| pDENV1-4 | YFV | 66.5 | 51.8 | 4.2 | 15.5 |
| pZIKV | ⌀ | 12.6 | 29.7 | 4.2 | 16.6 |
| pZIKV | D1-4 | 78.7 | 51.2 | 32.5 | 30.9 |
| pZIKV | ZIKV | 88.4 | 25.2 | 107.2 | 35.5 |
| pZIKV | YFV | 11.6 | 33.0 | 4.9 | 14.8 |

**Repeat for 2nd NP color (i.e., blue)**
- Vary Ab1 (on nitrocellulose
- Vary Ab 2 (conjugated to nanoparticle)
- Run dipsticks with NS1 for DENV1-4, ZIKV, and YFV and blank

- Image tests for grayscale intensity at test area

- Generate matrix of test line intensities for all combinations

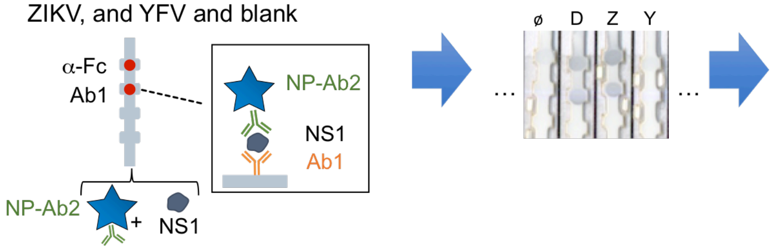


| | | Blue Nstar | | | |
|---|---|---|---|---|---|
| | | pDENV1-4 | | pZIKV | |
| Nitrocellulose | | test | control | test | control |
| pDENV1-4 | ⌀ | 5.1 | 22.9 | 6.6 | 15.6 |
| pDENV1-4 | D1-4 | 75.7 | 69.1 | 43.4 | 51.0 |
| pDENV1-4 | ZIKV | 43.3 | 66.8 | 44.3 | 64.5 |
| pDENV1-4 | YFV | 37.1 | 54.5 | -0.6 | 13.5 |
| pZIKV | ⌀ | 4.0 | 17.3 | 3.6 | 19.1 |
| pZIKV | D1-4 | 36.8 | 55.8 | 13.6 | 31.8 |
| pZIKV | ZIKV | 92.3 | 69.1 | 83.1 | 79.8 |
| pZIKV | YFV | 5.7 | 18.7 | 1.8 | 13.8 |

**Figure S4. Procedure for screening antibody pairs.**

### Color deconvolution

For pseudocolor identification, we used the image analysis method of color deconvolution (or "stain deconvolution") described by Ruifrok et al. 2001. It is a method used in histology where stains have overlapping RGB values, so when two stains are spatially colocalized, using RGB values is not able to distinguish between the stains. This results in a loss of information (i.e., one cannot accurately assign a pixel to belong to one stain or another). Color deconvolution is an orthonormal transformation of the RGB data into "stain channels" (called pseudocolors here) for each stain (here, the nanoparticles).

This allows us to find the pseudocolors of red nanospheres (NPs) and blue nanostars (GNS) when dried at the nitrocellulose. Using the pseudocolors, we can quantify the contribution of each nanoparticle when both are captured at the same spot. In short, this method uses the optical density to determine the "concentration" of NPs and GNS captured on the nitrocellulose when run together against the pathogens. The optical density is directly proportional to the NP concentration and can be calculated by the measured RGB intensities following this equation:

$$\boldsymbol{OD} = -\log_{\mathbf{10}}\left(\frac{\boldsymbol{I}}{\boldsymbol{I_0}}\right) = \boldsymbol{M} \cdot \boldsymbol{C}$$

Where: I is the measured RGB intensity, $I_0$ is the maximum intensity in the gray scale, C is what they call the stain concentration (NP in our case), M is the absorption factor and OD is the optical density.

The absorption factors (M) represents the contribution of each RGB channel in the OD space for each species (NP and GNS) run alone, and it is measured as one of the eigenvectors. The other one corresponds to the background of the picture. Thus, the absorption factor of the red NPs were measured in the OD space when the NPs were captured at the nitrocellulose. The same process was repeated with an image of GNS run alone in the test.

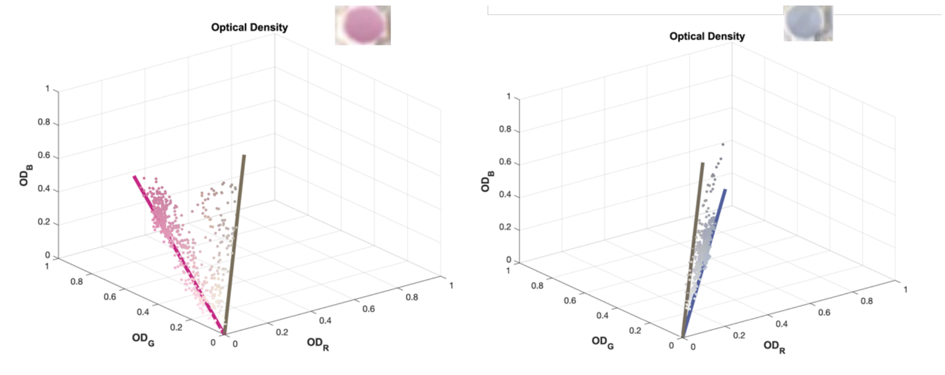


**Figure S5. OD of each the red NPs (left) and blue GNS (right) run alone.**

Knowing the absorption factors of the NPs and GNS (i.e. pseudocolors), one can determine which is the concentration of these NPs when captured together in the same nitrocellulose area. This method defines a third absorption factor for the background as the vectorial cross product of the two previous absorption factors. When NPs and GNS were run together against each pathogen, they may generate a sandwich immunoassay where both NPs are captured at different concentrations. This method allowed us to determine the amount of each nanoparticle captured at the test line by knowing the contribution of the pure NPs and GNS in each RGB channel (i.e. the pseudocolor).

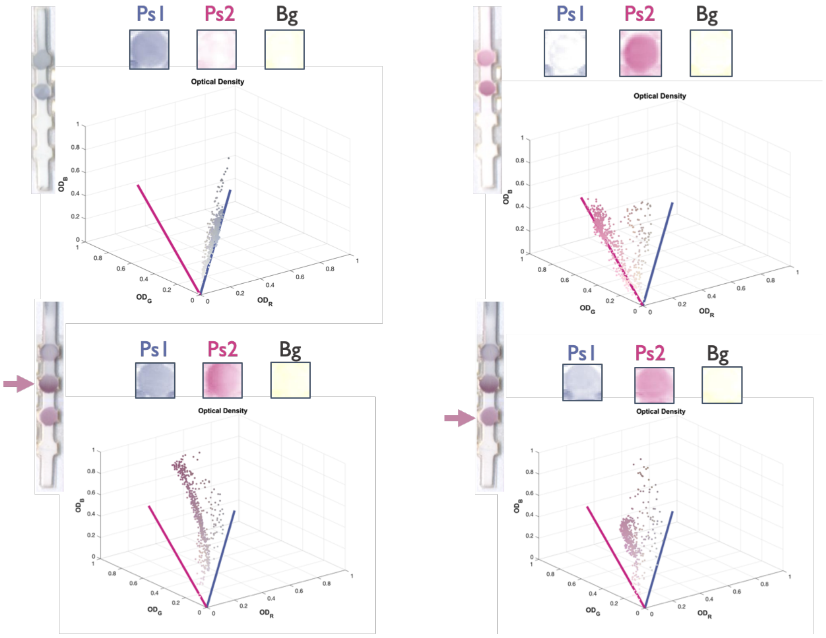


**Figure S6. Upper: individual nanoparticles and pseudocolor determination (left, GNS, right, NPs). Lower: Mixtures of the nanoparticles. The test spot (arrow) can be deconvolved into the contribution of the blue GNS (Ps1) and NPs (Ps2) and background (Bg). Two different deconvolutions are shown.**

The advantage of using this method is that we are not assuming that the blue intensity read from the RGB channels is solely proportional to the blue GNS concentration only, as the red NPs also contribute to blue intensity. Thus, when the PCA model was fed with the concentration of the NPs, GNS, and background, we improved the clusterization of the classes, as we were feeding the exact concentration of red NPs and blue GNS retained at the nitrocellulose area.

Ruifrok AC, Johnston DA. Quantification of histological staining by color deconvolution. *Anal Quant Cytol Histol* 23: 291-299, 2001.
